# Anoxic age as a new tool to predict biogeochemical consequences of oxygen depletion in lakes

**DOI:** 10.1101/2022.07.26.501517

**Authors:** Richard LaBrie, Michael Hupfer, Maximilian P. Lau

## Abstract

Lake deoxygenation is of growing concern because it threatens ecosystem services delivery. Complete deoxygenation, anoxia, is projected to prolong and expand in lakes, promoting the production or release of nutrients, greenhouse gases and metals from water column and the sediments. Accumulation of these compounds cannot be easily predicted thus hindering our capacity to forecast the ecological consequences of global changes on aquatic ecosystems. Here, we used lakes Arendsee and Mendota monitoring data to develop a novel metric, anoxic age, characterizing lake hypolimnetic anoxia. Anoxic age explained, as a single predictor, 44% to 58% of the variation for ammonium, soluble reactive phosphorus and a dissolved organic matter fluorophore. Anoxic age could be modelled using only two oxygen profiles and lake bathymetry, making it an easily applicable tool to interpret and extrapolate biogeochemical data. This novel metric thus has the potential to transform widely available oxygen profiles into an ecologically meaningful variable.

**Scientific Significance Statement:** Oxygen depletion in deep water layers of lakes is of growing concern as it expands due to eutrophication and climate change. Anoxia is deleterious to benthic invertebrates and fishes, enables the production of potent greenhouse gases and releases stored phosphorus from sediments, among others. However, quantitatively forecasting the consequences of anoxia remains a challenge. Here, we developed a novel metric, anoxic age, which may be derived from oxygen profiles to predict end-of-summer concentration of various water chemical parameters. We argue that all by-products of anaerobic microbial metabolism should be related to anoxic age as they are released or processed continuously during anoxia. We believe that anoxic age can be used to predict the ecological consequences of temporally and spatially growing anoxia.

## Introduction

Lakes provide essential ecosystem services (Jane et al. 2021), several of which are threatened by anthropic activities. Both eutrophication and global warming critically affect dissolved oxygen (DO) availability in lakes through higher hypolimnetic oxygen demand (Müller et al. 2012) or reduced hypolimnetic ventilation (Bartosiewicz et al. 2019). These pressures thus threaten more lake hypolimnia to become or stay anoxic for longer temporal episodes (Jenny et al. 2016; Matzinger et al. 2010). DO depletion in lake hypolimnia have far-reaching ecological consequences, including the accumulation of reduced compounds toxic to organisms, loss of habitats and intensified production of greenhouse gases (Jane et al. 2021, and references therein). Anoxia is also associated with phosphorus release from redox-sensitive sediment components (Hupfer and Lewandowski 2008). Monitoring and forecasting of these threats can be improved through modelling of oxygen dynamics.

Modelling approaches commonly differentiate between DO consuming processes in hypolimnetic sediments and waters (Livingstone and Imboden 1996), where contribution of the latter may be negligible in oxygen models for deep and clear lakes (Matzinger et al. 2010). These DO consumption models rely on widely available oxygen data and lake bathymetry and are a convenient tool to study onset and extent of anoxia, but not its consequences. Approaches to estimate consequences of anoxia, e.g., phosphorus release, used the lake-scale proportion of either sediment surface (Nürnberg 1984) or water volume (Foley et al. 2012) affected by anoxia, but both are insufficient to deconstruct the temporal sequence of anoxia-related processes. Currently, no modelling approach includes all elements required to fully address lake anoxia, including time, benthic and pelagic prokaryotic activity and their associated metabolite dynamics in the water column.

Lake water column chemistry reflects material take-up and release patterns of photosynthetic and heterotrophic organisms. The aphotic zone is dominated by heterotrophic processing of organic matter, consuming oxygen and releasing polyphenolic (Dadi et al. 2017) and fluorescent (FDOM, Burdige et al. 2004) dissolved organic matter (DOM) compounds. Under anoxic conditions, organic matter is not only the precursor for anaerobic metabolic products ammonium (NH_4_^+^) and methane, but also facilitates the production of oxygen-sensitive toxins as methylmercury and hydrogen sulfide (Achá et al. 2018), all accumulating in the hypolimnion during summertime anoxia. Thus, the extent and intensity of anoxia determines the quantity of these substances that will be introduced to surface waters in subsequent turnover events. However, easily predicting their accumulation remains a challenge.

In this study, we explored how to predict the accumulation of a common anaerobic metabolite, NH_4_^+^, soluble reactive phosphorus (SRP) and FDOM solely from lake oxygen data. To this end, we developed a novel metric, anoxic age, which characterizes how long discrete hypolimnion layers were DO depleted. Our underlying rational is that vertical exchange is considered negligible in stratified hypolimnetic waters (Rippey and McSorley 2009), and major solutes are either produced in situ or released from the conterminous sediments (Livingstone and Imboden 1996). With anoxic ages, we argue that not only the three compounds used herein, but all other reduced compounds produced as by-products of anaerobic metabolic pathways may be modelled with high precision and low effort.

To establish anoxic age as a widely applicable tool to study anoxia, we calculated anoxic ages from DO data acquired either in a small number of profiles or from several continuous loggers. We compared anoxic age values modelled from such widely available data formats to observed anoxic age values obtained in a specifically instrumented lake, running high-resolution DO profiler in (bi-)daily casts. We based all calculations on the Livingstone and Imboden (1996) deductive model between layer-specific oxygen consumption rates (J_z_) and sediment area to water volume ratio (α(z)). We kept the deductive approach but found that reconstructing daily hypolimnetic oxygen profiles was improved using non-linear equations, thus allowing for accurate prediction of anoxic age and hence of anaerobic metabolites, including NH_4_^+^, SRP and a FDOM component from limited data. Anoxic age thus proves to be an easily modelled metric describing anoxic lake biogeochemistry that can predict a wide array of compound accumulation and turnover.

## Materials and methods

### Study sites

We place our research in two eutrophic lakes that develop anoxia during summer stratification: Lake Arendsee (Germany) and Lake Mendota (USA) (table S1, Kreling et al. 2017; Ladwig et al. 2021). We used (bi-)daily multiparameter profiles (YSI) and 5 oxygen loggers (D-Opto, Zebra-Tech, New Zealand) data from the Lake Arendsee monitoring program (Hupfer et al. 2019), and weekly to fortnightly multiparameter profiles (YSI Exo2) from Lake Mendota in 2018 and 2020 (Magnuson et al. 2021).

### Anoxic age calculation

The anoxic age concept transforms oxygen data below a specified threshold into an information-bearing and ecologically meaningful variable. In essence, anoxic age reflects the time that passed since a parcel of water crossed the threshold (Fig. S1). Here, we used a conventional DO threshold for anoxia, 2 mg O_2_ L^−1^ (Jenny et al. 2016; Rabalais et al. 2010). Hypolimnetic waters have decreasing oxygen concentrations that are typically monitored in multiple depths, where each stratum may be considered as discrete because of negligible turbulent vertical diffusion (Rippey and McSorley 2009). The anoxic age increases for every consecutive timestep (Day_i_) a water stratum DO concentration (DO_z_) is below the chosen threshold (DO_threshold_) (eq. 1), where *i* and *n* are the first and last day of seasonal stratification, respectively.

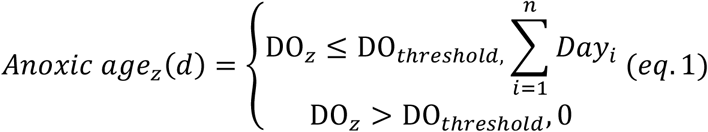

Anoxic age calculation requires DO concentration in high spatiotemporal resolution (e.g., daily and 1 m steps). Since measurements in this resolution are rarely available, oxygen values may be interpolated from two profiles, or alternatively derived from J_z_, themselves calculated from a small number of oxygen profiles (Livingstone and Imboden 1996). To assess if anoxic age may be accurately predicted from all common measurement practices, we subsampled observations from the full datasets of Lake Arendsee (see below).

### Morphometry

To calculate α(z), we used equations 2 and 3, where A(z) is the lake area at depth z, A_0_ is the lake surface area, z_m_ is the maximum depth of the lake and q is a fitting parameter (Livingstone and Imboden 1996).

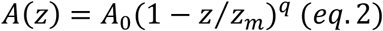

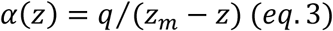

For both lakes, q was calculated using the hypsography. Lake Mendota hypsography is available on http://www.bathybase.org/Data/100-199/100/ (accessed 2021-10-18) and for Lake Arendsee, it was calculated following equation 5 in Håkanson (2005). Exponent q was then used to calculate α(z) (eq. 3).

### Modelling oxygen

Lakes are conveniently monitored in either several campaigns during the stratification season, or with a small number of depths-discrete loggers, enabling J_z_ calculation. Hence, we calculated J_z_ in two different ways. We compared J_z_ (slopes) of a linear regression of manually selected dates with a segmented regression using two breakpoints (Muggeo 2008) between DO concentration and time. Both methods yielded similar values (Fig. S2). We also calculated J_z_ for all profile casts taken 28 days apart when the lake was still in fully oxic condition as a “two oxic profiles” scenario, and for all three hypolimnetic DO loggers combinations from lake Arendsee. J_z_ for Lake Arendsee are reported in table S2 to S4 and in table S5 for Lake Mendota. We then compared three different types of equations to best describe the relationship between J_z_ and α(z). We used a linear fit, and to capture the asymptotic behavior, used log-linear and exponential-plateau fits (eq. 4), where b (non-negative) and k (0 ≤ k ≤ 15) are fitting parameters, and J_z,max_ is the maximum J_z_ fitted as a random parameter by the nlsLM function (Elzhov et al. 2016). The best model was chosen using R^2^, lowest root-mean-square error (RMSE) and Akaike Information Criterion.

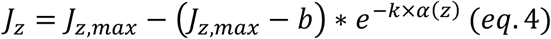

Using these equations, we modelled DO profiles series by assuming that the entire water column was fully oxygenated at the onset of summer stratification. Because stratification onset date is unknown, we needed to calibrate dates. To do so, we assigned dates to the DO profile series by matching a measured profile using lowest RMSE. For Lake Arendsee, we had measurements in sufficient temporal resolution to also calculate anoxic age directly from observations (AnoxA_obs_), but only used modelled anoxic ages (AnoxA_mod_) for Lake Mendota. We calculated the anoxic ages (eq. 1) and compared the first date (as day of year, DOY) on which each stratum became anoxic with observed first day of anoxia to assess quality of AnoxA_mod_.

### Chemical data

Concentrations of SRP and NH_4_^+^ were determined photometrically by molybdenum blue (Murphy and Riley 1962) and indophenol (Bolleter et al. 1961) methods, respectively, using segmented flow analysis (Scan++, Skalar Analytical, Netherlands). SRP and NH_4_^+^ values taken at non-integer depths were rounded at the nearest integer, and those at 0.5 m increment were floored as anoxia develops toward the surface. Exo2 FDOM was measured at excitation wavelength 365 nm, emission wavelength 480 nm and is expressed in quinine sulfate units (QSU). This peak, here referred to as F_365/480_ (F_ex/em_), is usually interpreted to indicate terrestrially derived recalcitrant compounds (peak C, Coble 1996).

All reported R^2^ were statistically significant at p-value < 0.01. All statistics and modelling were performed using R version 3.6.2 (R Core Team 2017) and all scripts and data will be available on github.com/MaxLauLab/AnoxicAge and attributed a doi using Zenodo.

## Results

### Nutrients and FDOM

We analyzed patterns in SRP and NH_4_^+^ from hypolimnetic waters in relation to anoxic age (AnoxA). When considering anoxic waters (AnoxA>0) only, we found a good relationship with SRP (R^2^ = 0.48), and with NH_4_^+^ (R^2^ = 0.44), where the slopes of these relationships (mean ± standard error), 0.73 ± 0.11 µg SRP L^−1^ d^−1^ and 6.0 ± 1.0 µg NH_4_^+^ L^−1^ d^−1^, represent whole anaerobic hypolimnion metabolism. Including AnoxA values of 0 (oxic waters) improved the relationship for both SRP (R^2^ = 0.62) and NH_4_^+^ (R^2^ = 0.55) but the rates then no longer represent anaerobic ecosystem-scale metabolism. In both cases, the relationships with DOY or depth alone were considerably weaker, and when combined explained nearly as much variation as AnoxA alone, suggesting that AnoxA captures all predictive power of these variables, but adding an ecologically meaningful interpretation. In multiple regressions, adding depth or DOY to AnoxA only slightly increased R^2^_adj_.

We explored the effect of anoxia on the fluorescent component F_365/480_ in lake Mendota using AnoxA_mod_. We found that AnoxA_mod_ explained a considerable fraction of F_365/480_ variation (Fig. 2b and d) with R^2^ of 0.63 in 2018 and R^2^ of 0.71 in 2020, when all data were considered. When only anoxic values were considered, R^2^ were lower at R^2^ = 0.56 and 0.58, respectively. Anoxia seemed to have a reproducible, stabilizing influence on this fluorescent component for 30 to 40 days (Fig. 2b and d); a behavior not well reflected by linear models and only visualized using AnoxA. In contrast to nutrients, the relationships with time (Fig. 2a and c) were better, with R^2^ of 0.84 in 2018 and 0.78 in 2020, respectively. Although time seemed to be a good predictor, its predictive power was greatly reduced when only values from anoxic waters were analyzed with R^2^ dropping to values of 0.32 to 0.27, respectively, indicating that the change in DOM composition is not constant over time spent in anoxia.

**Figure 1.**
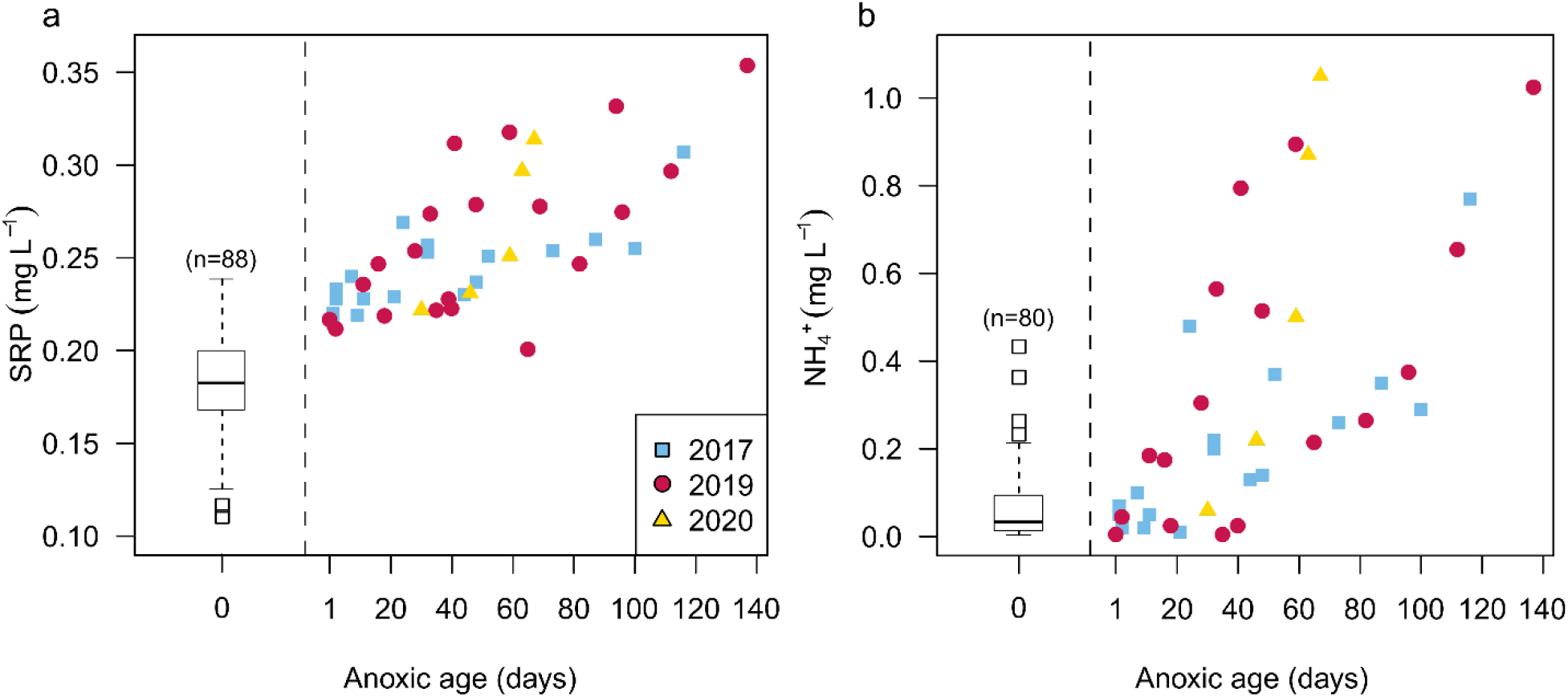
Soluble reactive phosphorus (SRP, a) and ammonium (NH_4_^+^, b) as a function of anoxic age in lake Arendsee. All SRP and NH_4_^+^ values at an anoxic age of 0 are displayed as a boxplot, with sample number above. Colored symbols represent different years; blue square: 2017, red circle: 2019 and yellow triangle: 2020.

**Figure 2.**
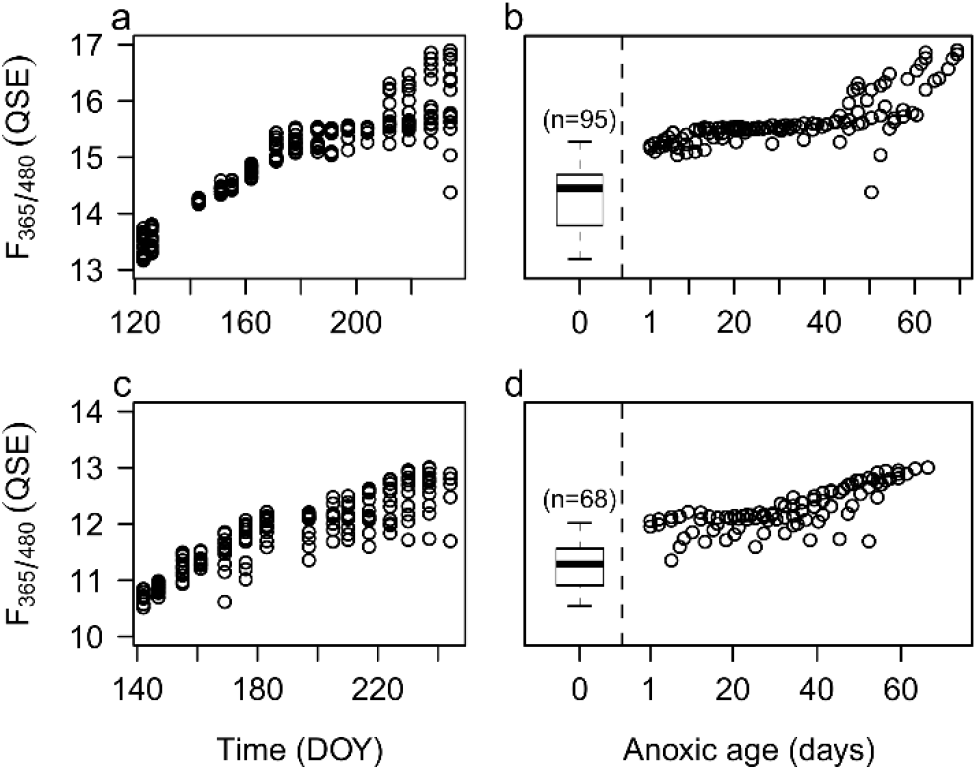
Fluorescent component F_365/480_ as a function of time (a, c) and modelled anoxic age (b, d), for 2018 (top row) and 2020 (bottom row) in lake Mendota.

### Modelling anoxic age

As AnoxA is calculated from daily DO profiles, we first assessed which type of equation best fitted J_z_ to α(z) from modelled profile time series. This step is critical to interpolate between irregular profile measurements or between missing depths of discrete loggers. The best fit was generally the exponential-plateau (Figures 3 and S3, table S6). When excluding several deep layers by subsampling J_z_ to simulate partial anoxia, we found that the exponential-plateau model was slightly less resilient than the log-linear relationship (Fig. S4). Overall, the linear fit was inferior to the other equations in both studied lakes (Fig. 3). The five loggers had lower fits regardless of equations, presumably because of a smaller number of α(z) values. Subsampling three loggers or using only two oxic profiles provided reliable J_z_-α(z) relationships in most cases (Fig. 4).

**Figure 3.**
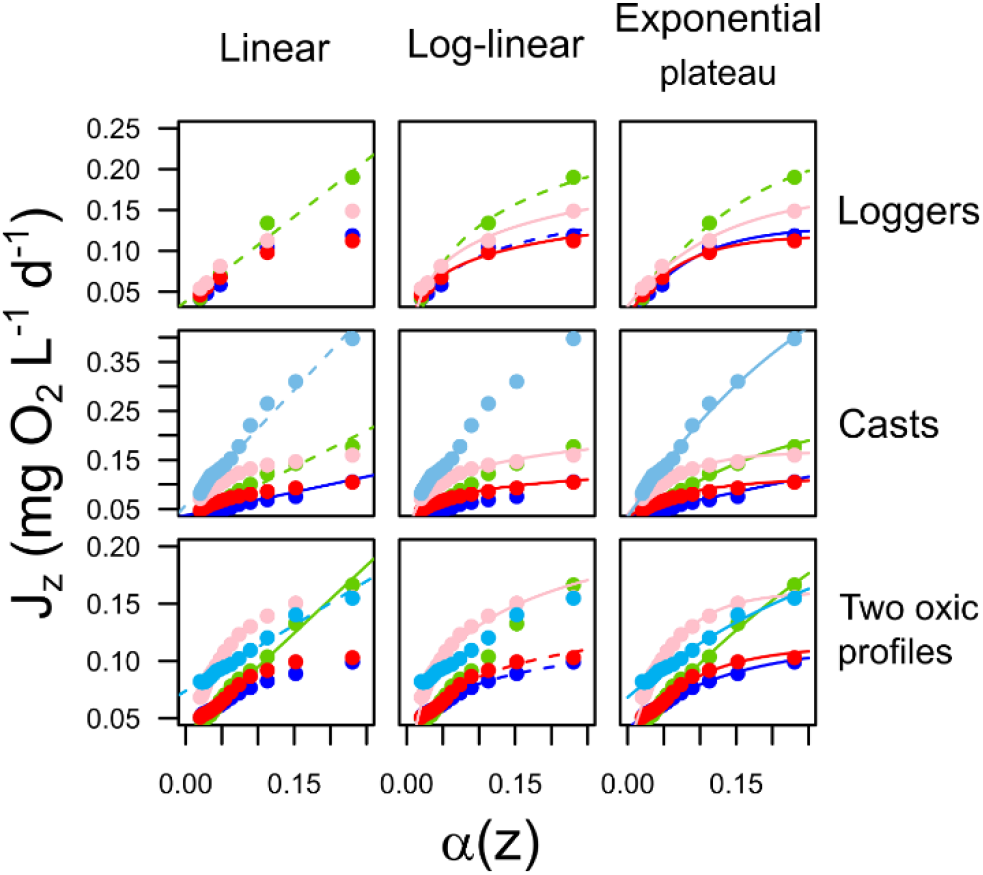
Oxygen consumption rates (J_z_) as a function of sediment area to volume ratio (α(z)) in Lake Arendsee. The rows represent autonomous loggers, autonomous (daily) YSI casts and the average of the two oxic profiles scenario, respectively. The columns represent the different fitting equations with linear, log-linear and exponential-plateau, respectively. Full lines in the different panels indicate fits between J_z_ and α(z) with R^2^ > 0.99, dashed lines, R^2^ > 0.97. Colors represent different years and are the same for all panels: blue (2017); green (2018); red (2019); pink (2020); light blue (2021).

**Figure 4.**
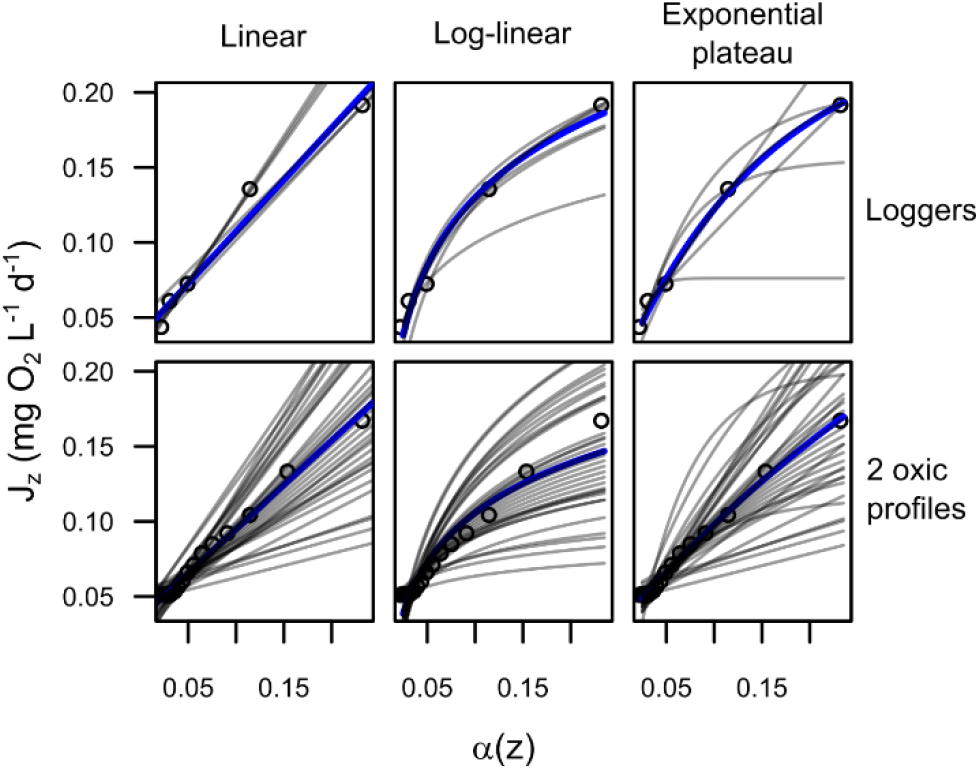
Oxygen consumption rates (J_z_) as a function of sediment area to volume ratio (α(z)) in Lake Arendsee by subsampling loggers (three, top row) and profiles casts (two, bottom row) to emulate common sampling scenarios (grey lines). The blue line represents the relationship using five loggers (top) and all profile casts (bottom).

Using linear, log-linear and exponential-plateau fits, we then compared their estimations of the date of anoxia’s first appearance. For all years and most data source-model type pairs, the log-linear and exponential-plateau models yielded good results, whereas the linear relationship did not (year 2019 in Fig. 5, other years in figures S5 to S8).

**Figure 5.**
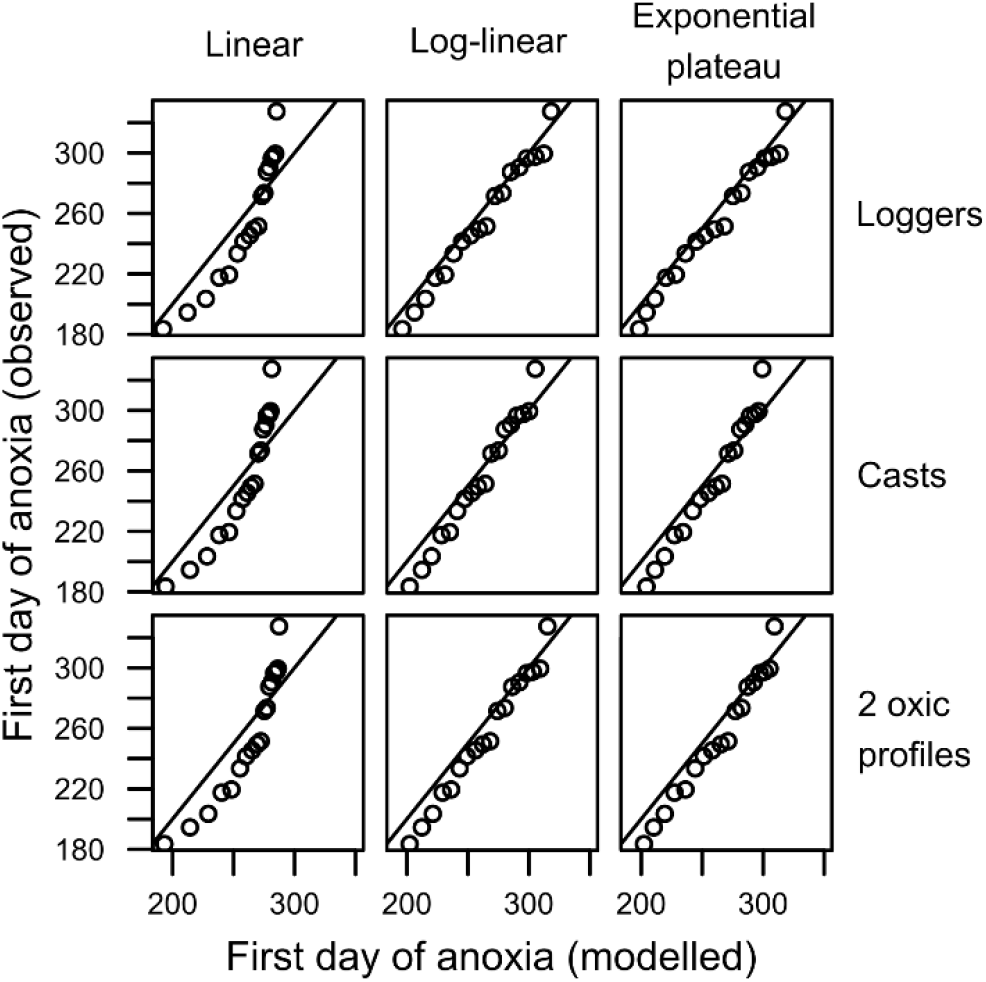
Relationship between observed and modelled first day of strata-specific anoxia in Arendsee, 2019. Each panel is a combination of data source (rows; loggers, autonomous YSI casts and two profiles during oxic conditions) and J_z_ model type (columns; linear, log-linear and exponential-plateau). Each point is a different hypolimnion depth (1 m increment), and the solid line represents a 1:1 relationship.

## Discussion

Water column and sediments heterotrophic respiration and oxidation of reduced compounds drive oxygen loss in lake hypolimnia (Matzinger et al. 2010; Steinsberger et al. 2020). To study and predict the specific lake water biogeochemistry that begins with deoxygenation, we developed a novel, ecologically meaningful metric, anoxic age. To this end, we built on the deductive approach to deconstruct oxygen depletion rates using lake bathymetry. In the original approach, fitting parameters of linear regressions directly represent sediments and water DO consumption rates (Livingstone and Imboden 1996; Rhodes et al. 2017). However, in both lakes studied herein, non-linear equations with a decreasing slope at higher α(z) values much better approximated the J_z_-α(z) relationships. These observations are consistent with other systems and can partly be explained by the smaller diffusive DO flux at smaller partial pressure gradients that dominate the deeper strata (Ladwig et al. 2021; Rippey and McSorley 2009). Non-linear fits lump areal and volumetric oxygen consumption rates but can predict more accurate daily oxygen profiles and consequently, anoxic age.

### Nutrients and FDOM

We examined the succession of anaerobic processes in lake hypolimnia over time and derived metabolic rates using linear models of substrate concentration over anoxic age. We estimated a lake’s anoxic ages based on a one-dimensional conceptualization of the water column, assuming that horizontal diffusivity greatly exceeds vertical diffusivity (Quay et al. 1980). Larger vertical diffusivity such as during seiching and horizontal turbulence promote vertical exchange among water layers, decreasing between-layer concentration differences (Rippey and McSorley 2009). Hence, the observed metabolic rates calculated herein would be slightly underestimated and therefore represent conservative estimates across all hypolimnetic layers.

Anoxic age is in essence a space-for-time transformation and examines the accumulation of anaerobic metabolic byproducts independent from the position in the water column. Therefore, we expect a stronger coupling of anoxic age with pelagic-driven anaerobic metabolites than with those related to sediment processes. The fluorescent component F_365/480_ that increased during Lake Mendota hypolimnetic anoxia is a dominant fluorescence signature in anoxic marine waters (Loginova et al. 2016) which suggests the production of recalcitrant DOC from both pelagic processing and sediment release (Dadi et al. 2017; Lau and del Giorgio 2020). Anoxic age revealed two stages of anoxic F_365/480_ transformation: a month-long stabilization followed by a strong increase of this component. This behavior remains unexplained by current conceptualization and would be missed without the anoxic age metric, offering a new lens to investigate carbon dynamics.

We also found good, positive relationships with SRP and NH_4_^+^, two compounds mostly related to sediment processes (Carey et al. 2022). The SRP increase likely reflected the initial phase of organic phosphorus diagenetic release, unobstructed by SRP-capturing iron oxides at the sediment surface (Hupfer and Lewandowski 2008). Classically, this process should be more pronounced in water layers with a higher α(z), yet we found a linear increase with depth-independent AnoxA, and models including depth as a variable were only slightly better. Similarly, anaerobic NH_4_^+^ production is dominated by sediment diagenesis, but also a product of pelagic microbial dissolved organic nitrogen turnover (Berman et al. 1999), and was adequately modeled with AnoxA alone. Limitations of molecular diffusion may be responsible for the depth-independent behavior of anoxic metabolites similar to oxygen consumption in sediments (Rippey and McSorley 2009). Therefore, a space-for-time analysis can capture the lake-scale trend of substrate behavior under anoxic conditions.

### Modelling oxygen profiles

We presented various equations to calculate anoxic age from widely available oxygen monitoring data. The exponential-plateau provided the best fit between J_z_ and α(z) for most lakes and years, and the best prediction of first day of anoxia. This test assumed a monotonic DO decrease, although physical mixing may oxygenate upper hypolimnetic strata even during stratification (Burns 1995), resulting in a reset of anoxic age to zero. This reset of anoxic age would be reflected in lake water biogeochemistry as reduced compounds are highly DO sensitive and rapidly oxidize (Kappler et al. 2004). This simplification seemed adequate for deep lakes’ hypolimnia but may need critical evaluation in shallower and more wind-exposed lakes. We note that an unconstrained exponential-plateau equation is particularly sensitive to J_z_ at large α(z). However, constraining the equation with *a priori* knowledge of the system provided adequate extrapolations, similar to the more robust log-linear equation. Thus, we recommend using the most plausible fit as the objective is to accurately predict anoxic ages.

The high temporal and spatial oxygen sampling resolution in Lake Arendsee enabled us to directly calculate anoxic ages, but also to test result quality of different sampling scenarios. By subsampling 3 DO loggers and by simulating two profiles per year, we assessed that these low vertical and temporal sampling resolutions were enough to adequately model oxygen consumption rates. With only two oxygen profiles, it is possible that deeper parts of the lake are already anoxic, prohibiting the use of these readings for J_z_ calculations. In this scenario, the linear and exponential-plateau were inadequate, but the log-linear equation nonetheless provided good J_z_-α(z) approximations. Common lake sampling practices therefore allow daily oxygen profiles modelling and thus anoxic age calculation.

### Future perspectives

Anoxia is pervasive and of growing concern in aquatic ecosystems worldwide (Rabalais et al. 2010), promoted by various anthropic activities including eutrophication and browning (Brothers et al. 2014; Jenny et al. 2016). We argue that anoxic age can be used across aquatic ecosystems to predict critical consequences of anoxia. Anoxic age can be tuned to specific DO_threshold_ of interest (table S7) to analyze the effects of anoxia on organisms (Elshout et al. 2013), greenhouse gases (Bastviken et al. 2002; Richardson et al. 2009) and toxic substances (Achá et al. 2018; Jorgensen et al. 1979; Sánchez-España et al. 2017). As lake-specific production rates can be modelled using anoxic age from limited observations, these production rates will provide valuable information to study drivers and trends of anaerobic metabolism and aid in assessing aquatic ecosystems health under global change.

## Data availability statement

All raw data and scripts will be available on Github.com/MaxLauLab/AnoxicAge upon manuscript acceptance and attributed a doi using zenodo.org.

## Acknowledgements

We would like to thank Sylvia Jordan (IGB) for managing the long-term program in Lake Arendsee and for her help with data validation. We are grateful to Christiane Herzog and Thomas Rossoll (IGB) for laboratory work and other technical support. We acknowledge Tobias Goldhammer (IGB) for discussions and for the support as head of the Chemical Laboratory. We’d like to thank Tom Shatwell (Helmholtz-Zentrum für Umweltforschung) for insightful discussions. The monitoring program is partly supported by the State Agency for Flood Protection and Water Management Saxony-Anhalt (LHW).

## Notes

http://www.bathybase.org/Data/100-199/100/

https://lter.limnology.wisc.edu/lake_mendota_multiparameter_sonde_profiles

### Competing Interest Statement

The authors have declared no competing interest.

## References

Achá, D. and others 2018. Algal Bloom Exacerbates Hydrogen Sulfide and Methylmercury Contamination in the Emblematic High-Altitude Lake Titicaca. Geosciences 8: 438.

Bartosiewicz, M., A. Przytulska, J. F. Lapierre, I. Laurion, M. F. Lehmann, and R. Maranger. 2019. Hot tops, cold bottoms: Synergistic climate warming and shielding effects increase carbon burial in lakes. Limnology and Oceanography Letters 4: 132–144.

Bastviken, D., J. Ejlertsson, and L. Tranvik. 2002. Measurement of Methane Oxidation in Lakes: A Comparison of Methods. Environmental Science & Technology 36: 3354–3361.

Berman, T., C. Béchemin, and S. Maestrini, Y. 1999. Release of ammonium and urea from dissolved organic nitrogen in aquatic ecosystems. Aquat. Microb. Ecol. 16: 295–302.

Bolleter, W. T., C. J. Bushman, and P. W. Tidwell. 1961. Spectrophotometric Determination of Ammonia as Indophenol. Analytical Chemistry 33: 592–594.

Brothers, S. and others 2014. A feedback loop links brownification and anoxia in a temperate, shallow lake. Limnol. Oceanogr. 59: 1388–1398.

Burdige, D. J., S. W. Kline, and W. Chen. 2004. Fluorescent dissolved organic matter in marine sediment pore waters. Mar. Chem. 89: 289–311.

Burns, N. M. 1995. Using hypolimnetic dissolved oxygen depletion rates for monitoring lakes. New Zealand journal of marine and freshwater research 29: 1–11.

Carey, C. C. and others 2022. Anoxia decreases the magnitude of the carbon, nitrogen, and phosphorus sink in freshwaters. Global Change Biology 28: 4861–4881.

Coble, P. G. 1996. Characterization of marine and terrestrial DOM in seawater using excitation-emission matrix spectroscopy. Mar. Chem. 51: 325–346.

Dadi, T., M. Harir, N. Hertkorn, M. Koschorreck, P. Schmitt-Kopplin, and P. Herzsprung. 2017. Redox Conditions Affect Dissolved Organic Carbon Quality in Stratified Freshwaters. Environmental Science & Technology 51: 13705–13713.

Elshout, P. M. F., L. M. Dionisio Pires, R. S. E. W. Leuven, S. E. Wendelaar Bonga, and A. J. Hendriks. 2013. Low oxygen tolerance of different life stages of temperate freshwater fish species. Journal of Fish Biology 83: 190–206.

Elzhov, T. V., K. M. Mullen, A.-N. Spiess, and B. Bolker. 2016. minpack.lm: R interface to the Levenberg-Marquardt nonlinear least-squares algorithm found in MINPACK, plus support for bounds. R package 1.2-1.

Foley, B., I. D. Jones, S. C. Maberly, and B. Rippey. 2012. Long-term changes in oxygen depletion in a small temperate lake: effects of climate change and eutrophication. Freshwater Biology 57: 278–289.

Håkanson, L. 2005. The Importance of Lake Morphometry for the Structureand Function of Lakes. International Review of Hydrobiology 90: 433–461.

Hupfer, M., A. Kleeberg, and J. Lewandowski. 2019. Internal pools and fluxes of phosphorus in dimictic lake Arendsee, Northeastern Germany, p. 169–185. In A. D. Steinman and B. Spears [eds.], Internal phosphorus loading in lakes: causes, case studies, and management. J. Ross Publishing.

Hupfer, M., and J. Lewandowski. 2008. Oxygen Controls the Phosphorus Release from Lake Sediments – a Long-Lasting Paradigm in Limnology. International Review of Hydrobiology 93: 415–432.

Jane, S. F. and others 2021. Widespread deoxygenation of temperate lakes. Nature 594: 66–70.

Jenny, J.-P. and others 2016. Urban point sources of nutrients were the leading cause for the historical spread of hypoxia across European lakes. Proceedings of the National Academy of Sciences 113: 12655.

Jorgensen, B. B., J. G. Kuenen, and Y. Cohen. 1979. Microbial transformations of sulfur compounds in a stratified lake (Solar Lake, Sinai)1. Limnol. Oceanogr. 24: 799–822.

Kappler, A., M. Benz, B. Schink, and A. Brune. 2004. Electron shuttling via humic acids in microbial iron(III) reduction in a freshwater sediment. FEMS Microbiology Ecology 47: 85–92.

Kreling, J., J. Bravidor, C. Engelhardt, M. Hupfer, M. Koschorreck, and A. Lorke. 2017. The importance of physical transport and oxygen consumption for the development of a metalimnetic oxygen minimum in a lake. Limnol. Oceanogr. 62: 348–363.

Ladwig, R. and others 2021. Lake thermal structure drives interannual variability in summer anoxia dynamics in a eutrophic lake over 37 years. Hydrology and Earth System Sciences 25: 1009–1032.

Lau, M. P., and P. del Giorgio. 2020. Reactivity, fate and functional roles of dissolved organic matter in anoxic inland waters. Biology Letters 16: 20190694.

Livingstone, D. M., and D. M. Imboden. 1996. The prediction of hypolimnetic oxygen profiles: a plea for a deductive approach. Can. J. Fish. Aquat. Sci. 53: 924–932.

Loginova, A. N., S. Thomsen, and A. Engel. 2016. Chromophoric and fluorescent dissolved organic matter in and above the oxygen minimum zone off Peru. Journal of Geophysical Research: Oceans 121: 7973–7990.

Magnuson, J. J., S. R. Carpenter, and E. H. Stanley. 2021. Lake Mendota Multiparameter Sonde Profiles: 2017 - current ver 2. Environmental Data Initiative.

Matzinger, A., B. Müller, P. Niederhauser, M. Schmid, and A. Wüest. 2010. Hypolimnetic oxygen consumption by sediment-based reduced substances in former eutrophic lakes. Limnol. Oceanogr. 55: 2073–2084.

Muggeo, V. 2008. Segmented: An R Package to Fit Regression Models With Broken-Line Relationships.

Müller, B., L. D. Bryant, A. Matzinger, and A. Wüest. 2012. Hypolimnetic Oxygen Depletion in Eutrophic Lakes. Environmental Science & Technology 46: 9964–9971.

Murphy, J., and J. P. Riley. 1962. A modified single solution method for the determination of phosphate in natural waters. Analytica Chimica Acta 27: 31–36.

Nürnberg, G. K. 1984. The prediction of internal phosphorus load in lakes with anoxic hypolimnia1. Limnol. Oceanogr. 29: 111–124.

Quay, P. D., W. S. Broecker, R. H. Hesslein, and D. W. Schindler. 1980. Vertical diffusion rates determined by tritium tracer experiments in the thermocline and hypolimnion of two lakes1,2. Limnol. Oceanogr. 25: 201–218.

R Core Team. 2017. R: A language and environment for statistical computing. Vienna, Austria: R Foundation for Statistical Computing. URL https://www.R-project.org/.

Rabalais, N. N., R. J. Díaz, L. A. Levin, R. E. Turner, D. Gilbert, and J. Zhang. 2010. Dynamics and distribution of natural and human-caused hypoxia. Biogeosciences 7: 585–619.

Rhodes, J., H. Hetzenauer, M. A. Frassl, K.-O. Rothhaupt, and K. Rinke. 2017. Long-term development of hypolimnetic oxygen depletion rates in the large Lake Constance. Ambio 46: 554–565.

Richardson, D., H. Felgate, N. Watmough, A. Thomson, and E. Baggs. 2009. Mitigating release of the potent greenhouse gas N2O from the nitrogen cycle – could enzymic regulation hold the key? Trends in Biotechnology 27: 388–397.

Rippey, B., and C. McSorley. 2009. Oxygen depletion in lake hypolimnia. Limnol. Oceanogr. 54: 905–916.

Sánchez-España, J. and others 2017. Anthropogenic and climatic factors enhancing hypolimnetic anoxia in a temperate mountain lake. Journal of Hydrology 555: 832–850.

Steinsberger, T., R. Schwefel, A. Wüest, and B. Müller. 2020. Hypolimnetic oxygen depletion rates in deep lakes: Effects of trophic state and organic matter accumulation. Limnol. Oceanogr.

