## Supplementary information for "Anoxic age as a new tool to predict biogeochemical consequences of oxygen depletion in lakes"

Richard LaBrie[1], Michael Hupfer[2] and Maximilian P. Lau[1, 2]

[1] Interdisciplinary Environmental Research Centre, TU Bergakademie Freiberg, Akademiestraße 6, 09599 Freiberg, Germany

[2] Leibniz Institute of Freshwater Ecology and Inland Fisheries (IGB), Müggelseedamm 310, 12587 Berlin, Germany

### Additional figures and tables

Table S1. Study sites basic morphometry

| Study site | Max depth (m) | Surface area ($km^2$) | Coordinates | Loggers depth (m) |
|---|---|---|---|---|
| **Lake Arendsee** | 49 | 5.14 | 53°53’21’’N 11°28’27’’E | 30, 35, 40, 45, 47 |
| **Lake Mendota** | 25 | 39.4 | 43°06’24’’N 89°25’29’’W | - |

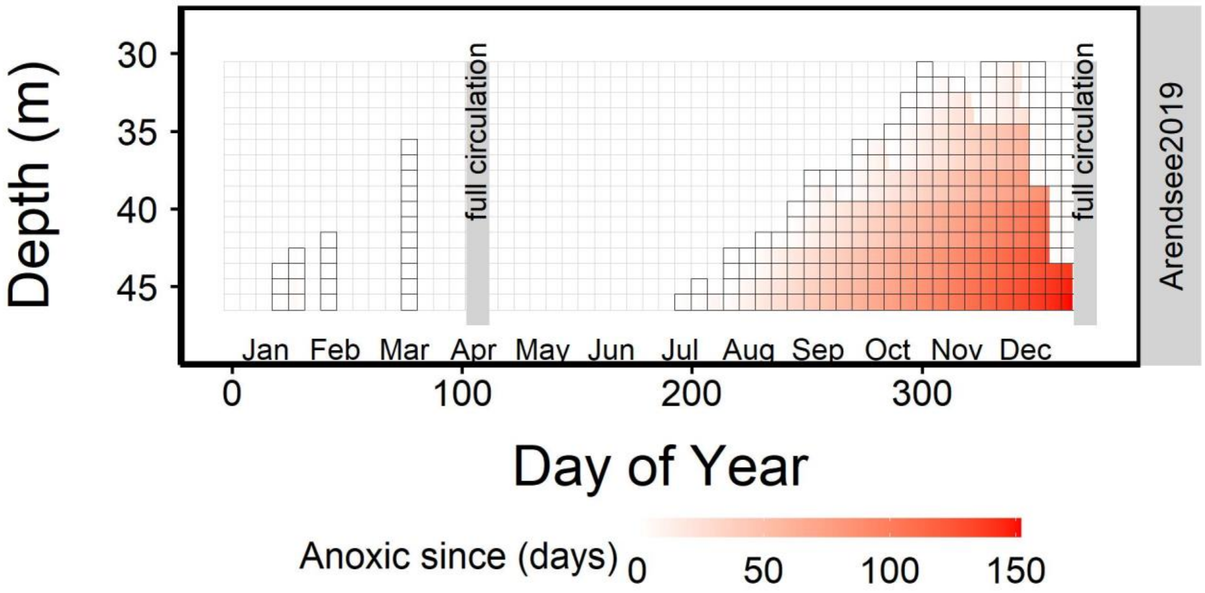


Figure S1. Schematic representation of anoxic age in Lake Arendsee hypolimnion in 2019. Grey bars represent lake overturning in spring and winter. Colors represent anoxic age, where longer period spent in anoxia are darker colored.

Segmented regression divides dissolved oxygen (DO) timeseries into episodes of different DO consumption patterns, effectively separating episodes with little or no DO change (turnover, later-summer anoxia) from the continuous DO consumption during the summer. We see that both methods yield identical values at low $J_z$ values and that higher values tend to be higher (i.e., slope are steeper) using the value from the segmented regression method (Fig. S1).

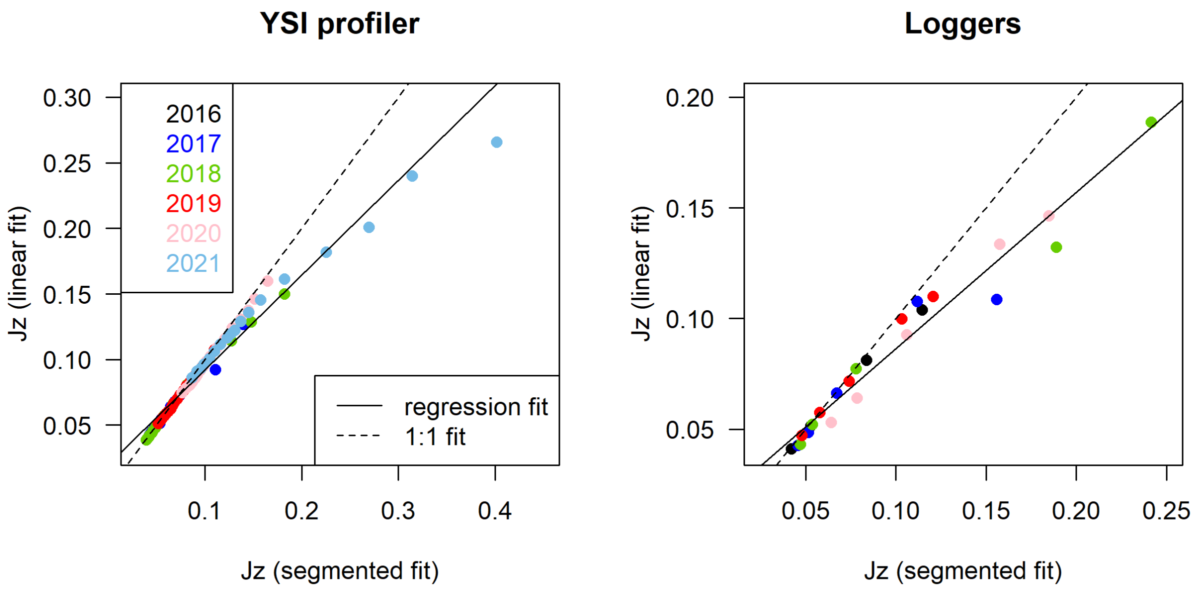


Figure S2. Comparison of linear oxygen consumption over times using manually selected dates and a segmented regressions with two breakpoints. We see that both methods yield identical values at low $J_z$ values and that higher values tend to be higher using the segmented regression method.

Table S2. $J_z$ values and standard error calculated from loggers in Lake Arendsee

| Depth | Year | $J_z$ | Error | Year | $J_z$ | Error | Year | $J_z$ | Error | Year | $J_z$ | Error | Year | $J_z$ | Error |
|---|---|---|---|---|---|---|---|---|---|---|---|---|---|---|---|
| **30** | | 0.0416 | 0.0004 | | 0.0444 | 0.0005 | | 0.0450 | 0.0008 | | 0.0489 | 0.0002 | | 0.0568 | 0.0006 |
| **35** | | 0.0508 | 0.0004 | | 0.0509 | 0.0006 | | 0.0623 | 0.0004 | | 0.0583 | 0.0003 | | 0.0637 | 0.0007 |
| **40** | 2016 | 0.0730 | 0.0008 | 2017 | 0.0614 | 0.0005 | 2018 | 0.0737 | 0.0007 | 2019 | 0.0702 | 0.0004 | 2020 | 0.0841 | 0.0013 |
| **45** | | 0.0841 | 0.0020 | | 0.1068 | 0.0013 | | 0.1369 | 0.0042 | | 0.1008 | 0.0013 | | 0.1154 | 0.0032 |
| **47** | | NA | NA | | 0.1217 | 0.0027 | | 0.1928 | 0.0046 | | 0.1151 | 0.0022 | | 0.1516 | 0.0033 |

Table S3. $J_z$ values and standard error calculated from profile casts in Lake Arendsee

| Depth | Year | $J_z$ | Error | Year | $J_z$ | Error | Year | $J_z$ | Error | Year | $J_z$ | Error | Year | $J_z$ | Error |
|---|---|---|---|---|---|---|---|---|---|---|---|---|---|---|---|
| **30** | | 0.0415 | 0.0007 | | 0.0391 | 0.0005 | | 0.0517 | 0.0005 | | 0.0761 | 0.0005 | | 0.0868 | 0.0009 |
| **31** | | 0.0430 | 0.0007 | | 0.0406 | 0.0006 | | 0.0537 | 0.0005 | | 0.0803 | 0.0006 | | 0.0916 | 0.0009 |
| **32** | | 0.0450 | 0.0007 | | 0.0416 | 0.0007 | | 0.0548 | 0.0005 | | 0.0833 | 0.0006 | | 0.0959 | 0.0009 |
| **33** | | 0.0465 | 0.0007 | | 0.0428 | 0.0007 | | 0.0564 | 0.0005 | | 0.0845 | 0.0006 | | 0.0989 | 0.0009 |
| **34** | | 0.0476 | 0.0007 | | 0.0447 | 0.0007 | | 0.0579 | 0.0005 | | 0.0867 | 0.0006 | | 0.1043 | 0.0010 |
| **35** | | 0.0481 | 0.0007 | | 0.0459 | 0.0008 | | 0.0596 | 0.0006 | | 0.0894 | 0.0007 | | 0.1098 | 0.0011 |
| **36** | | 0.0486 | 0.0007 | | 0.0484 | 0.0008 | | 0.0617 | 0.0006 | | 0.0917 | 0.0007 | | 0.1158 | 0.0012 |
| **37** | | 0.0496 | 0.0007 | | 0.0505 | 0.0009 | | 0.0644 | 0.0007 | | 0.0948 | 0.0007 | | 0.1230 | 0.0013 |
| **38** | | 0.0512 | 0.0007 | | 0.0526 | 0.0010 | | 0.0661 | 0.0007 | | 0.0977 | 0.0008 | | 0.1264 | 0.0013 |
| **39** | 2017 | 0.0530 | 0.0007 | 2018 | 0.0552 | 0.0012 | 2019 | 0.0681 | 0.0008 | 2020 | 0.1013 | 0.0009 | 2021 | 0.1308 | 0.0011 |
| **40** | | 0.0524 | 0.0008 | | 0.0579 | 0.0014 | | 0.0709 | 0.0008 | | 0.1057 | 0.0010 | | 0.1371 | 0.0010 |
| **41** | | 0.0546 | 0.0009 | | 0.0668 | 0.0014 | | 0.0740 | 0.0009 | | 0.1120 | 0.0012 | | 0.1450 | 0.0011 |
| **42** | | 0.0576 | 0.0010 | | 0.0814 | 0.0013 | | 0.0779 | 0.0009 | | 0.1202 | 0.0014 | | 0.1575 | 0.0013 |
| **43** | | 0.0642 | 0.0009 | | 0.0927 | 0.0015 | | 0.0810 | 0.0011 | | 0.1282 | 0.0015 | | 0.1824 | 0.0019 |
| **44** | | 0.0684 | 0.0010 | | 0.1053 | 0.0017 | | 0.0852 | 0.0012 | | 0.1367 | 0.0016 | | 0.2253 | 0.0028 |
| **45** | | 0.0735 | 0.0012 | | 0.1270 | 0.0022 | | 0.0907 | 0.0014 | | 0.1444 | 0.0018 | | 0.2698 | 0.0035 |
| **46** | | 0.0802 | 0.0015 | | 0.1476 | 0.0026 | | 0.0976 | 0.0016 | | 0.1515 | 0.0021 | | 0.3146 | 0.0047 |
| **47** | | 0.1104 | 0.0022 | | 0.1820 | 0.0037 | | 0.1095 | 0.0017 | | 0.1648 | 0.0025 | | 0.4020 | 0.0075 |
| **48** | | 0.1394 | 0.0023 | | - | - | | - | - | | - | - | | - | - |

Table S4. $J_z$ values and median absolute deviation (MAD) calculated from "two oxic profiles" scenario in Lake Arendsee. MAD was used in this case to provide a type of error similar to a Theil-Sen slope.

| Depth | Year | $J_z$ | Error | Year | $J_z$ | Error | Year | $J_z$ | Error | Year | $J_z$ | Error | Year | $J_z$ | Error |
|---|---|---|---|---|---|---|---|---|---|---|---|---|---|---|---|
| **30** | | 0.0491 | 0.0030 | | 0.0524 | 0.0011 | | 0.0529 | 0.0016 | | 0.0703 | 0.0011 | | 0.0841 | 0.0020 |
| **31** | | 0.0507 | 0.0032 | | 0.0525 | 0.0011 | | 0.0540 | 0.0017 | | 0.0709 | 0.0014 | | 0.0835 | 0.0019 |
| **32** | | 0.0521 | 0.0033 | | 0.0530 | 0.0009 | | 0.0547 | 0.0017 | | 0.0726 | 0.0014 | | 0.0833 | 0.0017 |
| **33** | | 0.0528 | 0.0033 | | 0.0527 | 0.0008 | | 0.0551 | 0.0016 | | 0.0758 | 0.0016 | | 0.0836 | 0.0017 |
| **34** | | 0.0532 | 0.0034 | | 0.0521 | 0.0008 | | 0.0560 | 0.0017 | | 0.0792 | 0.0018 | | 0.0837 | 0.0016 |
| **35** | | 0.0538 | 0.0034 | | 0.0522 | 0.0008 | | 0.0567 | 0.0018 | | 0.0830 | 0.0020 | | 0.0847 | 0.0017 |
| **36** | | 0.0547 | 0.0034 | | 0.0537 | 0.0008 | | 0.0574 | 0.0019 | | 0.0869 | 0.0020 | | 0.0864 | 0.0017 |
| **37** | | 0.0557 | 0.0035 | | 0.0548 | 0.0010 | | 0.0587 | 0.0020 | | 0.0905 | 0.0020 | | 0.0880 | 0.0019 |
| **38** | 2017 | 0.0566 | 0.0036 | 2018 | 0.0581 | 0.0012 | 2019 | 0.0603 | 0.0021 | 2020 | 0.0937 | 0.0020 | 2021 | 0.0905 | 0.0022 |
| **39** | | 0.0576 | 0.0038 | | 0.0615 | 0.0015 | | 0.0624 | 0.0023 | | 0.0982 | 0.0022 | | 0.0932 | 0.0029 |
| **40** | | 0.0593 | 0.0038 | | 0.0673 | 0.0019 | | 0.0650 | 0.0022 | | 0.1035 | 0.0024 | | 0.0948 | 0.0030 |
| **41** | | 0.0619 | 0.0041 | | 0.0725 | 0.0024 | | 0.0690 | 0.0022 | | 0.1103 | 0.0028 | | 0.0968 | 0.0032 |
| **42** | | 0.0652 | 0.0045 | | 0.0800 | 0.0029 | | 0.0748 | 0.0021 | | 0.1166 | 0.0029 | | 0.0991 | 0.0038 |
| **43** | | 0.0694 | 0.0047 | | 0.0862 | 0.0033 | | 0.0813 | 0.0020 | | 0.1251 | 0.0031 | | 0.1041 | 0.0048 |
| **44** | | 0.0741 | 0.0048 | | 0.0934 | 0.0035 | | 0.0886 | 0.0021 | | 0.1321 | 0.0033 | | 0.1114 | 0.0059 |
| **45** | | 0.0798 | 0.0051 | | 0.1057 | 0.0039 | | 0.0939 | 0.0023 | | 0.1413 | 0.0038 | | 0.1223 | 0.0075 |
| **46** | | 0.0862 | 0.0054 | | 0.1345 | 0.0059 | | 0.1013 | 0.0028 | | 0.1525 | 0.0053 | | 0.1423 | 0.0103 |
| **47** | | 0.0956 | 0.0067 | | 0.1685 | 0.0087 | | 0.1050 | 0.0037 | | 0.1574 | 0.0077 | | 0.1568 | 0.0139 |

Table S5. $J_z$ values and standard error calculated from profile casts in Lake Mendota

| Depth | Year | $J_z$ | Error | Year | $J_z$ | Error |
|---|---|---|---|---|---|---|
| **10** | 2018 | 0.159 | 0.004 | 2020 | 0.150 | 0.012 |
| **11** | | 0.163 | 0.004 | | 0.154 | 0.015 |
| **12** | | 0.167 | 0.005 | | 0.157 | 0.011 |
| **13** | | 0.173 | 0.005 | | 0.166 | 0.013 |
| **14** | | 0.175 | 0.005 | | 0.169 | 0.010 |
| **15** | | 0.177 | 0.005 | | 0.172 | 0.008 |
| **16** | | 0.183 | 0.005 | | 0.178 | 0.009 |
| **17** | | 0.191 | 0.007 | | 0.184 | 0.010 |
| **18** | | 0.200 | 0.010 | | 0.196 | 0.012 |
| **19** | | 0.209 | 0.009 | | 0.211 | 0.014 |
| **20** | | 0.214 | 0.009 | | 0.211 | 0.014 |
| **21** | | 0.216 | 0.009 | | - | - |
| **22** | | 0.218 | 0.010 | | - | - |
| **23** | | 0.221 | 0.010 | | - | - |

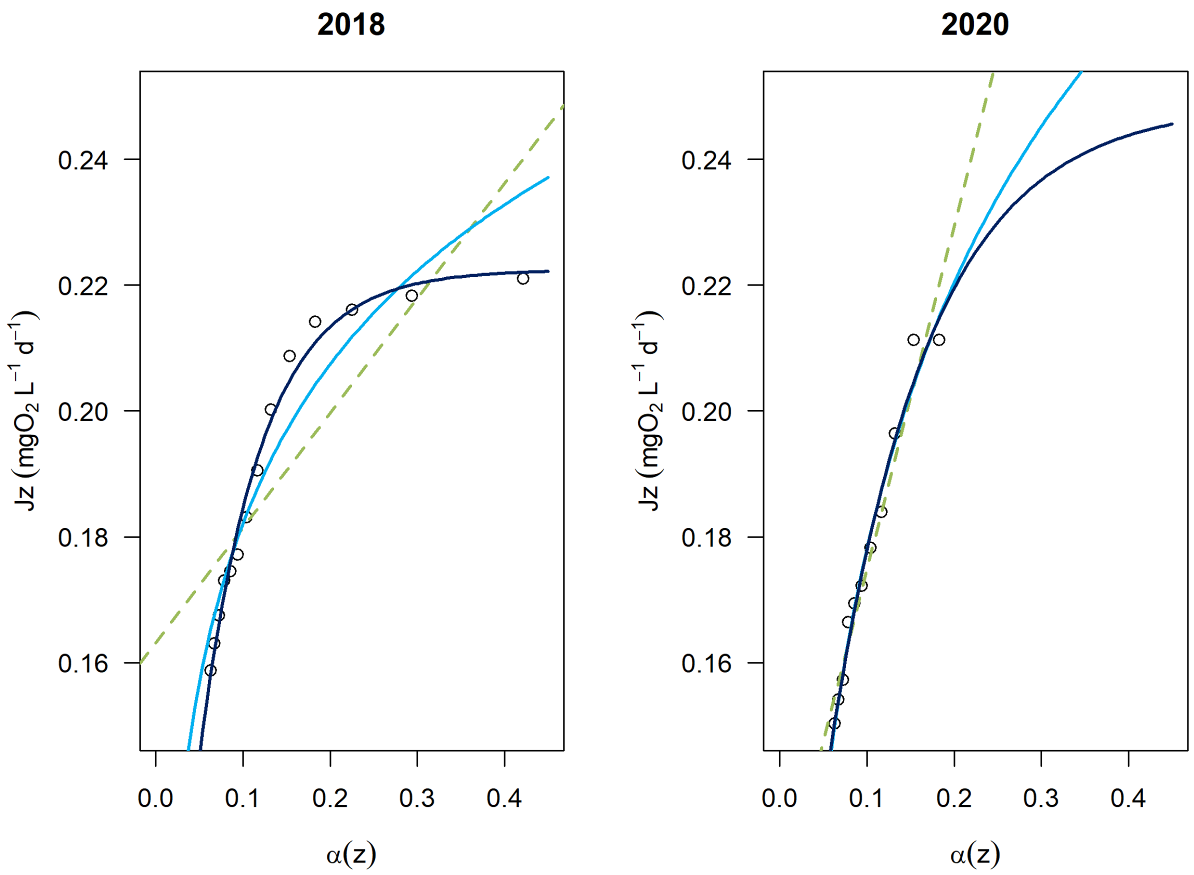


Figure S3. Oxygen consumption rates ($J_z$) as a function of sediment area to volume ratio ($\alpha(z)$) in lake Mendota. Green dashed-line: linear fit; light blue line: log-linear fit; dark blue line: exponential plateau fit.

Table S6. Summary table of $R^2$, residual mean square error (RMSE) and Akaike information criterion (AIC) of oxygen consumption ($J_z$) with sediment to volume ratio ($\alpha(z)$) relationships in lake Arendsee. Emphasis used to show the best model(s).

| | | Linear | | | Log-linear | | | Exponential plateau | | |
|---|---|---|---|---|---|---|---|---|---|---|
| Method | Year | $R^2$ | RMSE | AIC | $R^2$ | RMSE | AIC | $R^2$ | RMSE | AIC |
| Loggers | 2017 | 0.88 | 0.0106 | -25.3 | 0.97 | 0.0053 | -32.2 | **0.99** | **0.0037** | **-33.9** |
| | 2018 | 0.97 | 0.0097 | -26.1 | 0.98 | 0.0078 | -28.3 | **0.997** | **0.0032** | **-35.2** |
| | 2019 | 0.089 | 0.0085 | -27.4 | 0.99 | 0.0023 | -40.3 | **0.999** | **0.0008** | **-48.8** |
| | 2020 | 0.96 | 0.0073 | -29.0 | 0.99 | 0.0029 | -38.1 | **0.996** | **0.0022** | **-39.0** |
| Casts | 2017 | **0.99** | **0.0021** | **-146.5** | 0.88 | 0.0088 | -100.1 | **0.99** | **0.0021** | -144.4 |
| | 2018 | 0.97 | 0.0074 | -119.6 | 0.95 | 0.0088 | -113.3 | **0.99** | **0.0034** | **-145.7** |
| | 2019 | 0.90 | 0.0050 | -133.9 | **0.999** | **0.0004** | **-221.2** | 0.99 | 0.0012 | -180.8 |
| | 2020 | 0.86 | 0.0097 | -109.9 | 0.99 | 0.0023 | -160.9 | **0.997** | **0.0013** | **-179.1** |
| | 2021 | 0.98 | 0.0110 | -105.3 | 0.94 | 0.0212 | -87.7 | **0.99** | **0.0065** | **-122.2** |
| Two oxic profiles | 2017 | 0.95 | 0.0027 | -155.3 | 0.98 | 0.0017 | -171.5 | **0.996** | **0.0008** | **-199.8** |
| | 2018 | 0.95 | 0.0071 | -120.8 | 0.98 | 0.0042 | -139.8 | **0.996** | **0.0018** | **-168.0** |
| | 2019 | 0.87 | 0.0062 | -126.2 | 0.97 | 0.0029 | -153.5 | **0.99** | **0.0020** | **-165.5** |
| | 2020 | 0.82 | 0.0112 | -104.5 | 0.98 | 0.0033 | -143.2 | **0.999** | **0.0006** | **-206.7** |
| | 2021 | 0.98 | 0.0088 | -113.2 | 0.92 | 0.00180 | -87.6 | **0.99** | **0.0069** | **-120.3** |

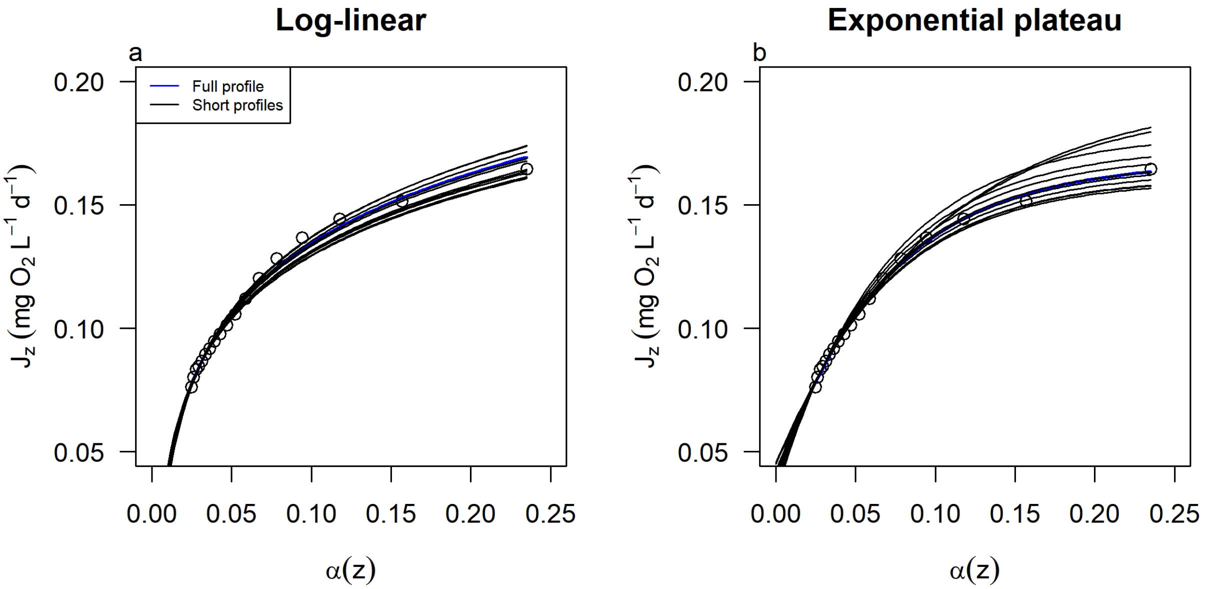


Figure S4. Oxygen consumption rates ($J_z$) as a function of sediment area to volume ratio ($\alpha(z)$) in lake Arendsee, 2020, using a log-linear equation (a) and an exponential plateau model (b). The blue line is the reference using the full dataset. The black lines represent the relationships between 5 to 15 subsampled $J_z$ and $\alpha(z)$starting from 30m and going down. This illustrates a second $O_2$ profile when the water column was already partially anoxic and $J_z$ could not be adequately calculated.

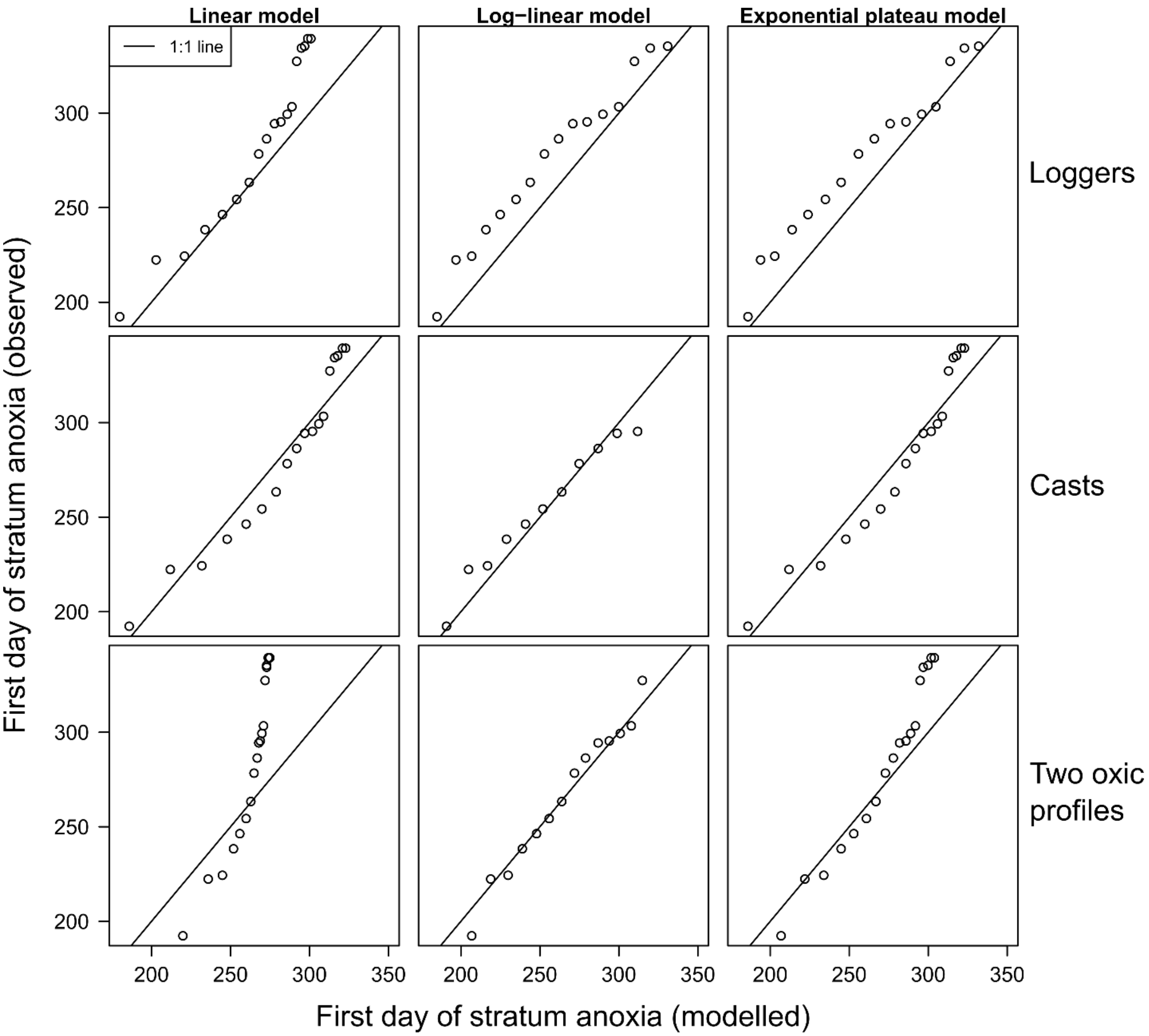


Figure S5. Relationship between observed and modelled first day of anoxia in Arendsee, 2017. Each panel is a combination of data source (rows; loggers, autonomous YSI casts and two oxic profiles) and model type (columns; linear, log-linear and exponential plateau). Each point is a different depth (1m increment) where the deepest depth is on the left, and the solid line represents a 1:1 relationship.

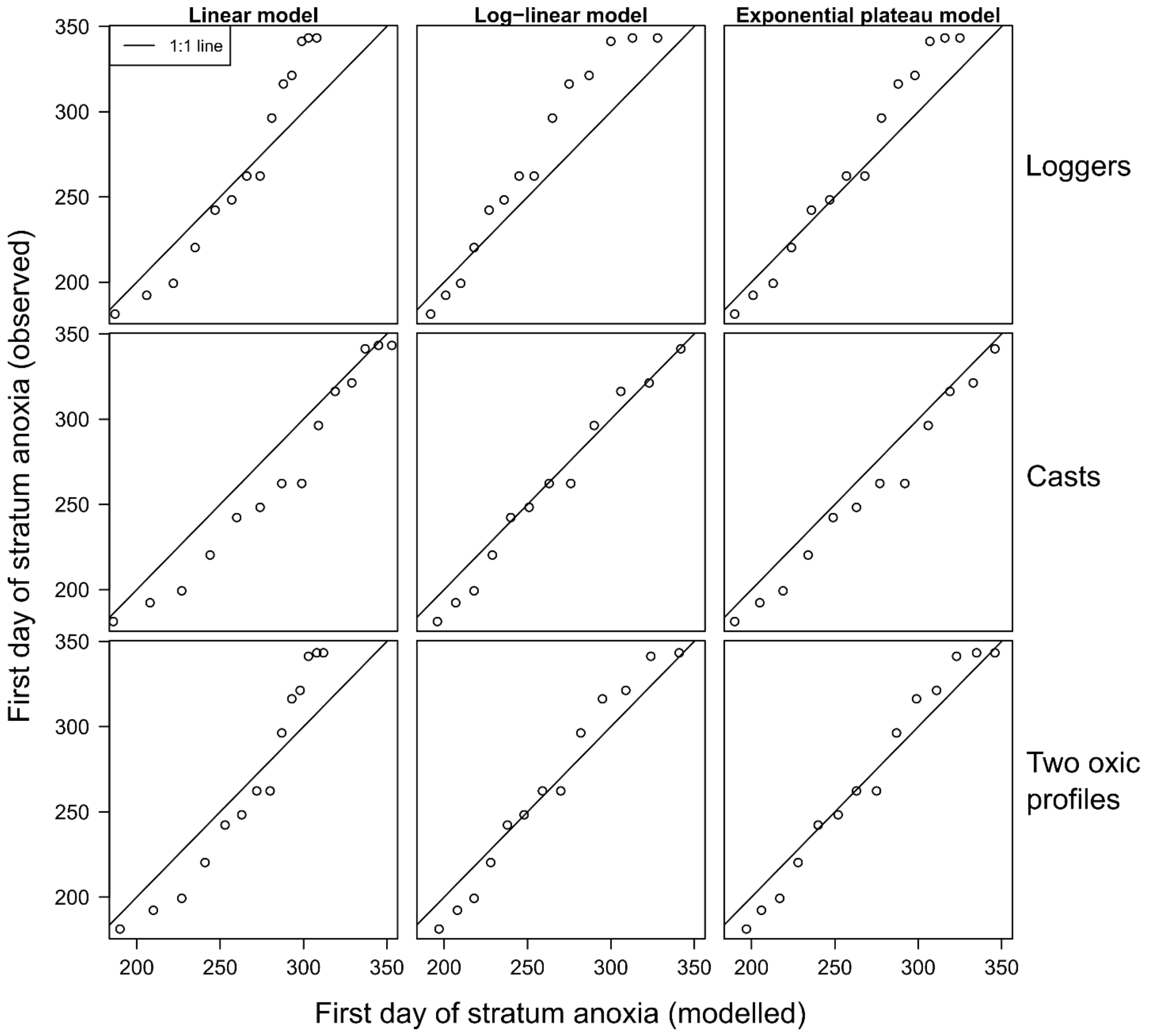


Figure S6. Relationship between observed and modelled first day of anoxia in Arendsee, 2018. Each panel is a combination of data source (rows; loggers, autonomous YSI casts and two oxic profiles) and model type (columns; linear, log-linear and exponential plateau). Each point is a different depth (1m increment) where the deepest depth is on the left, and the solid line represents a 1:1 relationship.

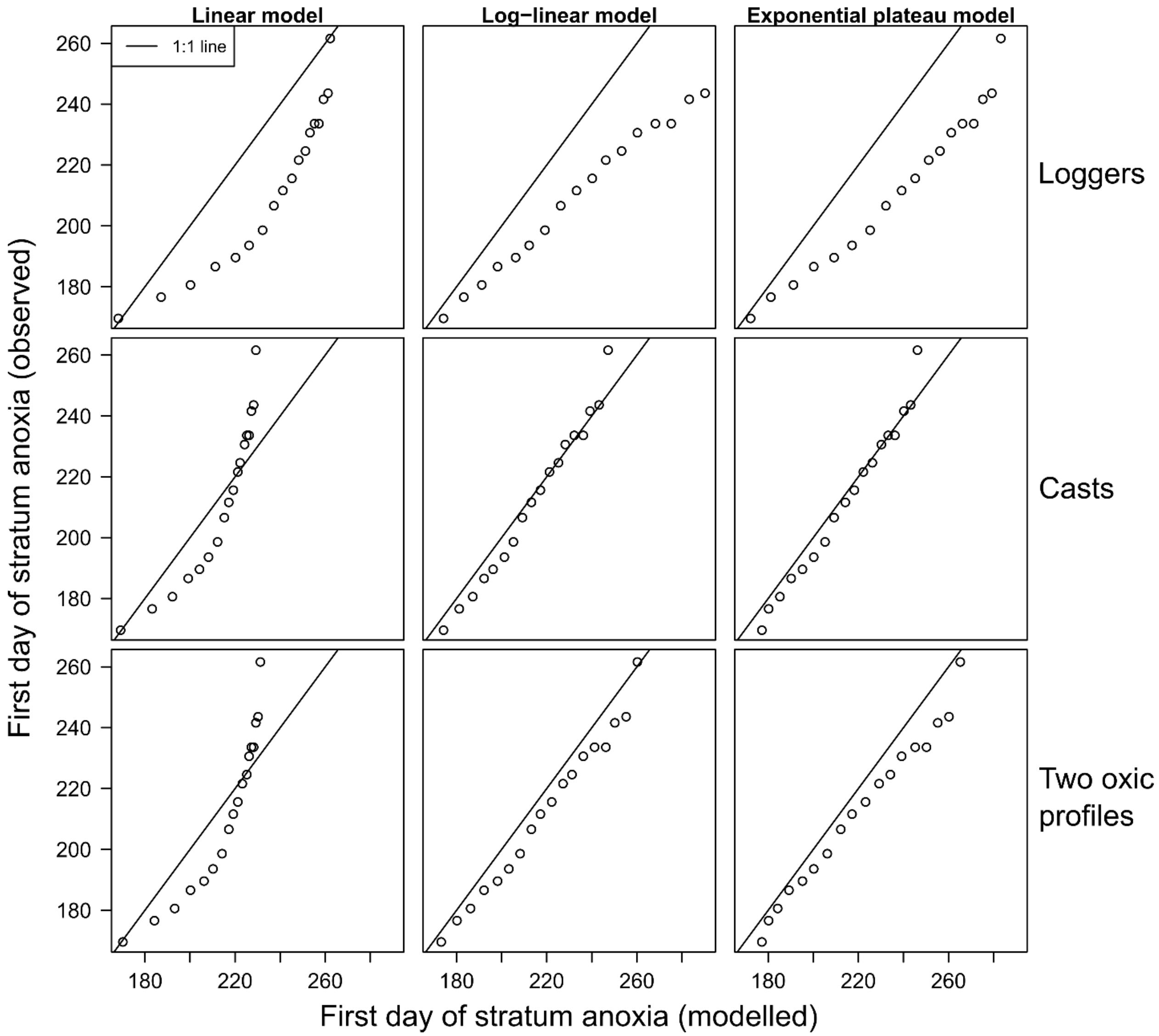


Figure S7. Relationship between observed and modelled first day of anoxia in Arendsee, 2020. Each panel is a combination of data source (rows; loggers, autonomous YSI casts and two oxic profiles) and model type (columns; linear, log-linear and exponential plateau). Each point is a different depth (1m increment) where the deepest depth is on the left, and the solid line represents a 1:1 relationship.

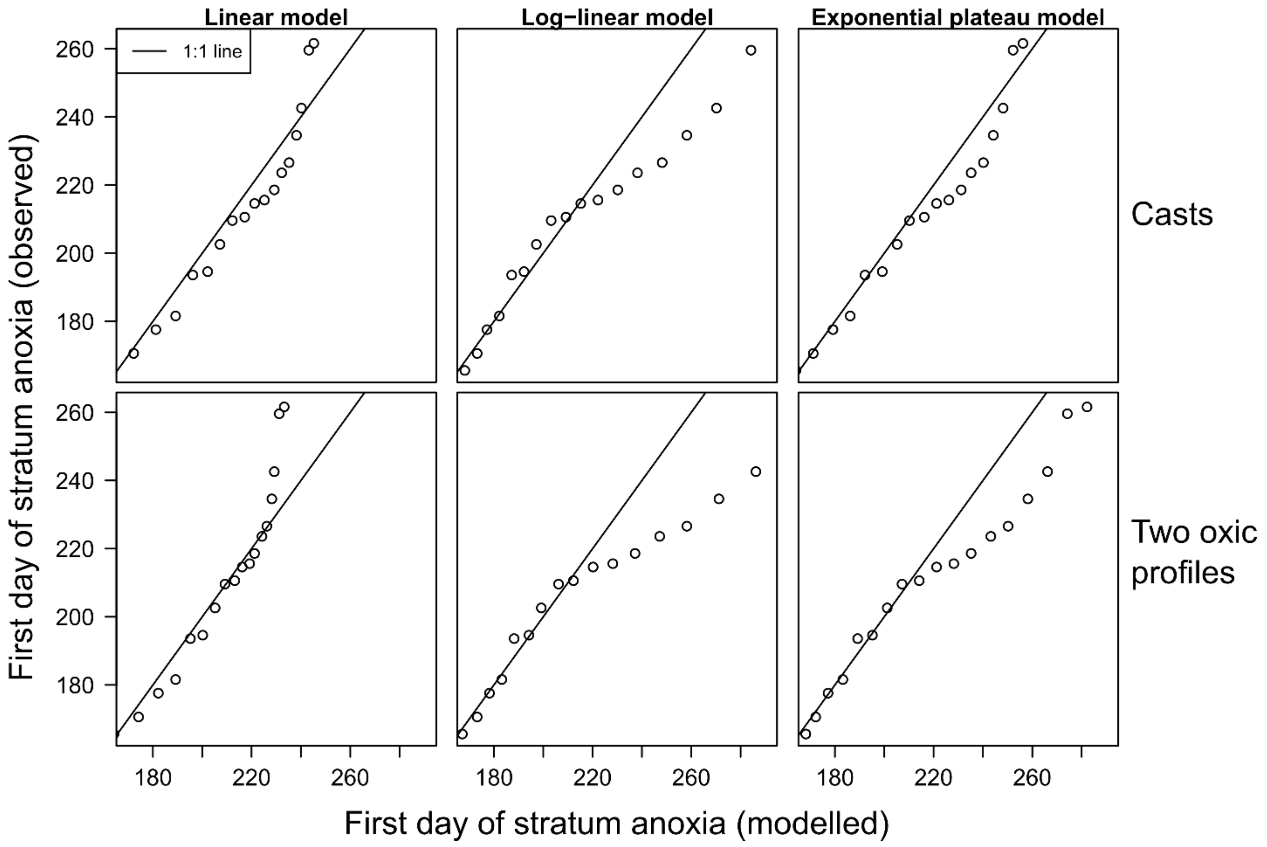


Figure S8. Relationship between observed and modelled first day of anoxia in Arendsee, 2021. Each panel is a combination of data source (rows; autonomous YSI casts and two oxic profiles) and model type (columns; linear, log-linear and exponential plateau). Each point is a different depth (1m increment) where the deepest depth is on the left, and the solid line represents a 1:1 relationship.

Table S7. Low oxygen threshold values for various organisms and reduced compounds. Reduced compounds contribute to hypolimnion deoxygenation before accumulating. $N_2O$ production is mostly stopped in anoxia where $N_2$ is preferentially produced. All references appear in the main text.

| | $O_2$ threshold (mg $L^{-1}$) | Impact | Note | Reference |
|---|---|---|---|---|
| Fish eggs | 3.1 | * | | Elshout et al. (2013) |
| Methane | ~ 0 | # | | Bastviken et al. (2002) |
| Nitrous oxide | $0 < O_2 \leq 2.5$ | # | | Richardson et al. (2009) |
| Metals | ~ 0 | # | Fe(II) and Mn(II)/(III) | Sánchez-España et al. (2017) |
| Hydrogen sulfide | ≤ 3.0<br>≤ 0.4 | # | High $SO_4^{2-}$ | Achá et al. (2018)<br>Jorgensen et al. (1979) |

* represents lethality, # represents water column accumulation.

### References in supplementary information


Achá, D. and others 2018. Algal Bloom Exacerbates Hydrogen Sulfide and Methylmercury Contamination in the Emblematic High-Altitude Lake Titicaca. Geosciences **8:** 438.

Bastviken, D., J. Ejlertsson, and L. Tranvik. 2002. Measurement of Methane Oxidation in Lakes: A Comparison of Methods. Environmental Science & Technology **36:** 3354-3361.

Elshout, P. M. F., L. M. Dionisio Pires, R. S. E. W. Leuven, S. E. Wendelaar Bonga, and A. J. Hendriks. 2013. Low oxygen tolerance of different life stages of temperate freshwater fish species. Journal of Fish Biology **83:** 190-206.

Jorgensen, B. B., J. G. Kuenen, and Y. Cohen. 1979. Microbial transformations of sulfur compounds in a stratified lake (Solar Lake, Sinai)1. Limnol. Oceanogr. **24:** 799-822.

Richardson, D., H. Felgate, N. Watmough, A. Thomson, and E. Baggs. 2009. Mitigating release of the potent greenhouse gas N2O from the nitrogen cycle – could enzymic regulation hold the key? Trends in Biotechnology **27:** 388-397.

Sánchez-España, J. and others 2017. Anthropogenic and climatic factors enhancing hypolimnetic anoxia in a temperate mountain lake. Journal of Hydrology **555:** 832-850.
